# Gender identification in Chicken (*Gallus gallus*) by PCR using whole blood and dried blood spot on filter paper as template: without prior DNA isolation

**DOI:** 10.1101/046888

**Authors:** S. Dhanasekaran, G. Dhinakar Raj, A. R. Vignesh, S. T. Selvan, B. Prakash, P. Perumal, Seenichamy Arivudainambi, Thambidurai Ganesh Babu

## Abstract

Accurate sex identification of pure line chickens in their early age has significant economic impact in breeding industry. In the recent years, range of Polymerase Chain Reaction (PCR) based sex determination techniques are routinely used to identify the sex of parent lines in breeding industries, however purified DNA is a prerequisite. Hence this study was aimed to develop a rapid and inexpensive PCR based gender identification method for chicken using whole blood samples and dried blood spots as template for PCR without DNA extraction. In addition, practicability of two W-chromosome specific gene targets in chicken for sex determination also characterised. Successful amplification of sex specific fragments and an internal control was achieved with the range of 0.125μl and 0.250μl volume of whole blood on filter paper (~1 mm) prepared from chicken and dried blood spot. This technique does not require DNA extraction, freeze/thawing of blood samples, pre-treatment with any reagents, dilution of whole blood or dried blood spots on filter paper. It can be carried out with commercially available Taq polymerase enzymes with increased concentration of MgCl_2_ (3 mM) and 0.5% of DMSO without optimisation of PCR buffers. In conclusion, as compared to the existing PCR based sex identification techniques, the present approach is relatively economic, time saving, requires minimal steps and eliminates the need for DNA extraction.

## Introduction

Sex determination in pure breed chickens in their early age is an important task to reduce the feeding cost by disposing unwanted sex, labour cost, achieve sex specific feeding programmes and commercialising the pure line birds based on their sex (Klein et al. 2003, Kaleta and Redmann 2008, Damme and Ristic, 2003). Gender in chicken and other avian species is being identified manually through cloacal or vent sexing, feather sexing by color of the plumage. Although manual gender determination techniques are inexpensive, it requires highly skilled personals and prone for more errors (Seemann, 2003). Other gender determination methods such as ultrasonography, flowcytometry, infrared spectroscopic imaging, laparoscopic method and molecular techniques such as cytogenetic method and hormonal assays requires highly trained personnel’s and sophisticated instrumentation facilities. Alternatively, PCR based techniques are used to determine the gender of avian species, common techniques like single-strand conformation polymorphism (SSCP), restriction fragment length polymorphism (RFLP), random amplification of polymorphic DNA (RAPD), amplification refractory mutation system (ARMS), sex-specific methylation pattern, SYBR green-based real time PCR combined with melting curve analysis and TaqMan probe based real time PCR are frequently used (Morinha et al. 2012, Caetano and Ramos. 2007, Rosenthal et al. 2010, Steiner et al. 2011, Chen et al. 2012, Granevitze et al. 2007, Ogawa et al. 1997).

Although PCR based sexing methods have several advantages over other sex determination techniques, it requires pure genomic DNA as template from biological specimens. For any genetic studies in avian species, DNA is being isolated from blastodermal cells, blood, feather barbs, muscle tissues, toe-pad skin, chorio-allantoic membrane and buccal swabs (Turkyilmaz et al 2010, Jensen et al. 2003, Trefil et al. 1999, Morinha et al. 2012, Rosenthal et al. 2010, Caetano and Ramos. 2007). Moreover, DNA isolation from blood or any other biological samples by various isolation procedures requires expensive reagents such as RNase A and proteinase-k and toxic reagents for instance phenol, chloroform, alcohol, also warrants the risk of cross contamination among individual sample. Therefore DNA isolation for large number of samples is not a promising approach in routine applications (Alonso, 2013).

Irrespective of biological specimens collected for DNA extraction to carryout molecular studies, the samples must be frozen or preserved immediately, before they begin to degrade. However dried blood spots (DBS) on filter paper or Guthrie cards do not require cold storage of dried blood samples. At present DBS cards are routinely used for prenatal genetic diagnosis, antibody detection, diagnosis of viral infections of humans or animals at field level (McCabe. 1991, McEwen JE and Reilly 1994, Parker and Cubitt, 1999). The practicability of dried blood spots on filter paper as specimen for sex determination of avian blood samples needs further evaluation, hence we also considered DBS on filter paper method in the present study.

As an alternative to the existing time consuming and expensive PCR based gender discrimination methods, we described the direct amplification of sex specific fragments from whole blood and dried blood spots on filter paper. In addition we also investigated the utility of two EST genes in sex determination of chicken.

## Materials and Methods

### Specimen collection and dried blood spot preparation

A total of 20 blood samples from young birds (male and female each n=10) of each species of chicken (*Gallus gallus*), Japanese quail (*Coturnix japonica*), duck (*Anas Platyrhynchos*) and turkey (*Meleagris gallopavo*) was collected from a small scale abattoir house located near Perambur (Corporation of Chennai), Chennai, Tamil Nadu, India. Prior to the collection of blood samples, the gender of each bird in each species was identified by morphology. Each blood sample was collected in equal amounts of triplicates separately with anti-coagulants of EDTA, sodium citrate and heparin. Five blood samples of Emu (*Dromaius novaehollandiae* and ostrich (*Struthio camelus*) was collected from Post Graduate Research Institute in Animal Sciences (Formerly known as Livestock Research Station), Tamil Nadu Veterinary and animal sciences University.

### Preparation of Dried blood spots

Dried blood spots on filter paper – 1 (Catalogue no: 1001125, GE Healthcare limited, UK) were prepared with 1-5 μl of blood by blotting within a dashed-line circles of 15 mm diameter, and the spots were dried 1-3 hrs under ambient temperature. After complete drying, the DBSs were stored in individual plastic bags at - 4°C. Dried blood spots were excised 1-2 mm diameter using sterile disposable biopsy punches for analysis and the remaining DBS cards sealed with individual cover and stored at 4°C for further use.

### DNA extraction from whole blood samples

DNA was extracted from blood samples of known male and female birds (each n=10) using Qiagen DNeasy Blood & Tissue Kit (Qiagen Inc, USA, Catalogue no: 69504) following manufacturer's protocol and dissolved in nuclease free water. DNA yield was determined by measuring the concentration of DNA in the eluate by its absorbance at 260 nm using BioPhotometer plus (Eppendorf, Germany).

### Primer designing and synthesis

Primer pairs specific for chicken W-chromosome specific genes and 18s rRNA were designed using PrimerSelect, DNA star software package (Primers listed in Table.1) synthesised by Eurofins India pvt Ltd (Bangalore, India). All the primer sequences were subjected to homology search using BLAST search against Chicken Genome in NCBI (National Center for Biotechnology Information).

### Polymerase Chain Reaction

All the PCR reactions were prepared in PCR workstation (Ultra-Violet Products Ltd, Cambridge, UK). An aliquot of blood sample (0.125 μl to 10 μl) or 1mm punch of dried blood spot on filter paper was added into individual 0.2-ml thin walled reaction tubes containing 5μl of 10X PCR buffer (100 mM Tris-HCl, pH 8.3, 500 mM KCl, 15 mM MgCl2 and 0.01% gelatin (Sigma Aldrich, Catalog No. P2192), 3 μl of 25mM MgCl_2_ (PCR grade, Sigma Aldrich, Catalog No. M8787), 0.5% of DMSO and 36μl of nuclease free water (Ambion^®^xs Nuclease-Free, Applied Biosystems, USA). The reaction was prepared at room temperature and mixed gently by pipetting to obtain a uniform solution, following incubation at 98°C for 3 min, the PCR tubes were briefly centrifuged. To the clear supernatant, 1μl of dNTP (0.2 mM each, Invitrogen, Catalog No. 18427) 5 units of Taq DNA polymerase for direct amplification of blood or DBS card and 2 units of enzyme for purified DNA amplification (Sigma Aldrich, Catalog No. D6677) and 1μl of 200 ?M each forward and reverse primer (primer sequence listed in Table 1) was added. Purified DNA from blood was used as a template in positive controls and nuclease free water was used as a negative control.

PCR reaction with direct blood as template was carried out with the following conditions, 95°C for 5 min, 35 cycles of denaturation at 95°C for 15sec, annealing for 30 sec at 54°C and extension at 72°C for 45 sec, followed by final extension at 72°C for 10 min.

PCR reaction with DBS punches was carried out with the following conditions, 98°C for 5 min, 37 cycles of denaturation at 95°C for 15sec, annealing for 30 sec at various temperature, listed in table 1, and extension at 72°C for 40 sec, followed by final extension at 72°C for 10 min. The PCR products were analysed in 2 % agarose gel and stained with ethidium bromide.

## Results

### W Chromosome specific gene targets of chickens

In the present study, chicken sex was determined using two W chromosome linked targets selected from previously reported cDNA micro array 2d-1H5 (Acc. No: AB188532) and 2d-2D9 (Acc. No: AB188526). These genes were localised in chicken W chromosome through *in silico* analysis and primer pairs was designed for these w-chromosome specific targets and 18s rRNA gene as an internal control to develop a duplex PCR assay (Primers and product size listed in Supplementary Table 1). In the Primer-BLAST search against chicken genome database, both the sex specific primer pairs had shown unique homology (100%) to W chromosome of chicken. Further bioinformatics analysis of primer targets for 2d-1H5 and 2d-2D9 had showed high similarity to two different regions of zinc finger SWIM domain-containing protein 6-like protein (Acc. No: NC006126).

### Sex determination of chicken by PCR using W chromosome specific primers

The ability of chicken W chromosome specific primers in gender differentiation was initially tested with DNA isolated from known male and female chicken blood samples. Both CHW1 FP&RP and CHW2 FP&RP have always produced unique female specific amplification in female chicken DNA samples. No amplification was observed in male DNA samples. Further a duplex PCR method was developed to determine the gender of chicken with primers for W chromosome specific (either CHW1 FP&RP or CHW2 FP&RP) and an endogenous control (18s rRNA) with different amplicon size (refer table 1). In the duplex PCR, female sex was determined by the presence of female specific 500bp (CHW1FP/1RP) along with 301 bp (CLD 18s 1FP/1RP) (Fig. 1) or 395 bp (CHW2FP/2RP) along with 195 bp (CLD 18s 2FP/2RP). While the male gender was identified only with only endogenous PCR products after agarose gel electrophoresis. The feasibility of these two sex specific primers in other commercially important avian species such as turkey, duck, quail, emu was carried out with DNA extracted from each species. CHW1FP/1RP primer pairs resulted in no amplification in both male and female DNA samples at various annealing temperature and MgCl_2_ concentration, whereas CHW2FP/RP primer sets produced similar non-specific amplification in both male and female DNA samples (result not shown).

**Fig. 1.**
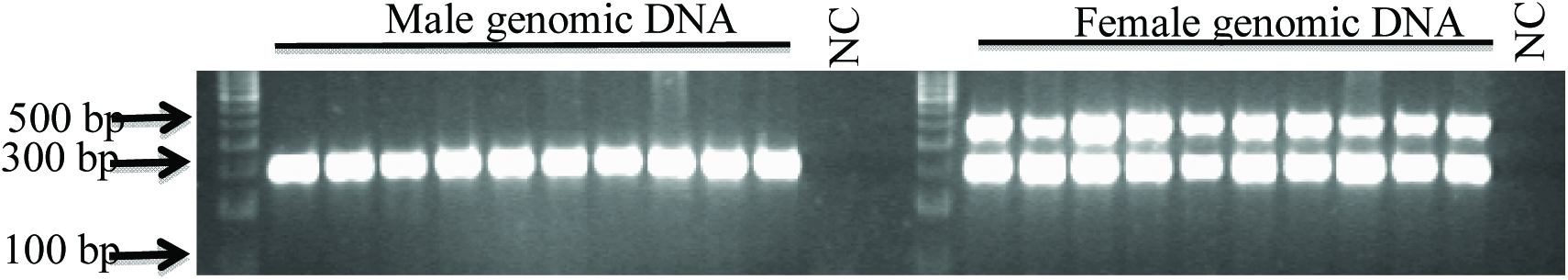
Gender identification of chicken (Gallus gallus) by PCR amplification of W chromosome specific sequences from genomic DNA isolated from whole blood. PCR was performed according to standard protocols, with genomic DNA isolated from male birds (n=10) and female birds (n=10) as the source of template. Taq DNA polymerase (1.5 U) and 200 ?M of each primer were added to a total reaction volume of 50 μl. Negative control was included in each reaction without the genomic DNA.

### Sexing of chicken using whole blood as a template

In our next approach, we studied the amplification of both sex specific and endogenous control target by duplex PCR using 1μl of chicken blood as template with commercially available PCR buffer and reagents without any modification. In our initial attempts, addition of 1μl whole blood samples to the PCR reaction failed to amplify many blood samples and the amplification intensity of certain samples is not satisfactory. Hence we tried with different volumes of whole blood of chicken ranging from 0.125μl to 10μl (fig 4). We observed that 0.125μl and 0.250μl of whole blood resulted reproducible and satisfactory PCR amplification of both sex specific and endogenous control. Also we observed that, use of 5 units of Taq polymerase enzyme in 50 μl PCR reactions instead of 2 units had resulted a satisfactory amplification of both the targets. We observed two distinct PCR products specific for CHW1FP/1RP primers (female specific 500 bp) and CLD 18s 1FP/1RP primer (internal control gene specific 300 bp) (Fig. 2). Whereas CHW2FP/2RP and CLD 18s 2FP/2RP primer pairs amplified, female specific 395 bp and an internal control specific 195 bp product (Fig. 5). The CHW2FP/2RP and CLD 18s 2FP/2RP combination produced a non-specific amplicon of around 800 bp, the preferred PCR products amplification intensity was not satisfactory.

**Fig. 2.**
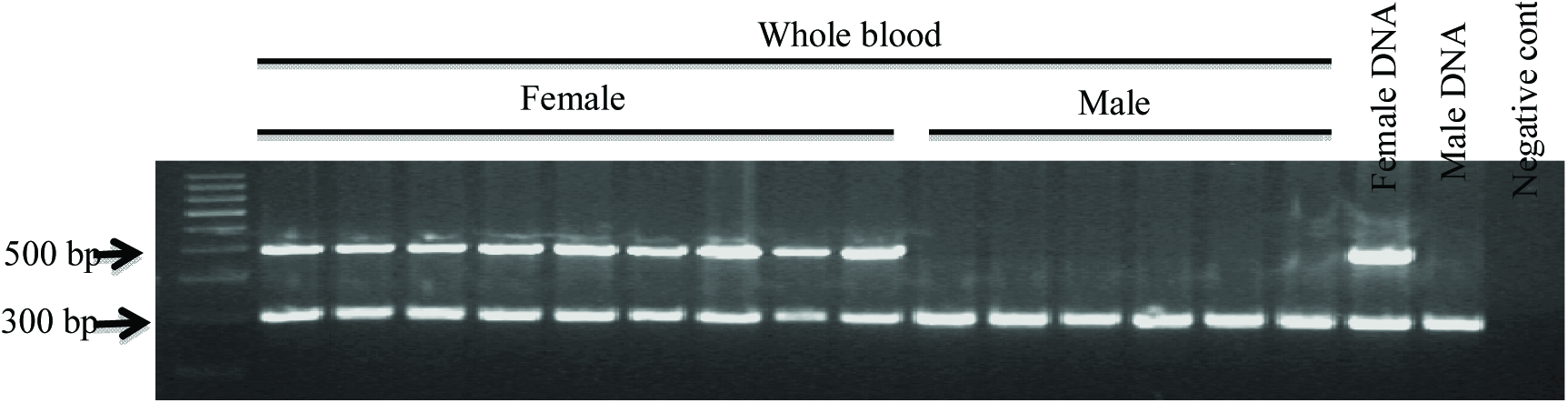
Gender identification of chicken (Gallus gallus) by PCR amplification of CHW1 sequences from whole blood (0.25μl). PCR was performed as described in the materials and methods using the whole blood from individual male birds (n=6) and female birds (n=9) as the source of template. Taq DNA polymerase (5 U) and 1μl of 200 ?M each forward and reverse primer were added to a total reaction volume of 50 μl. Genomic Dna isolated from male and female birds was used as positive control. Negative control was included in each reaction without the genomic DNA.

**Fig. 5.**
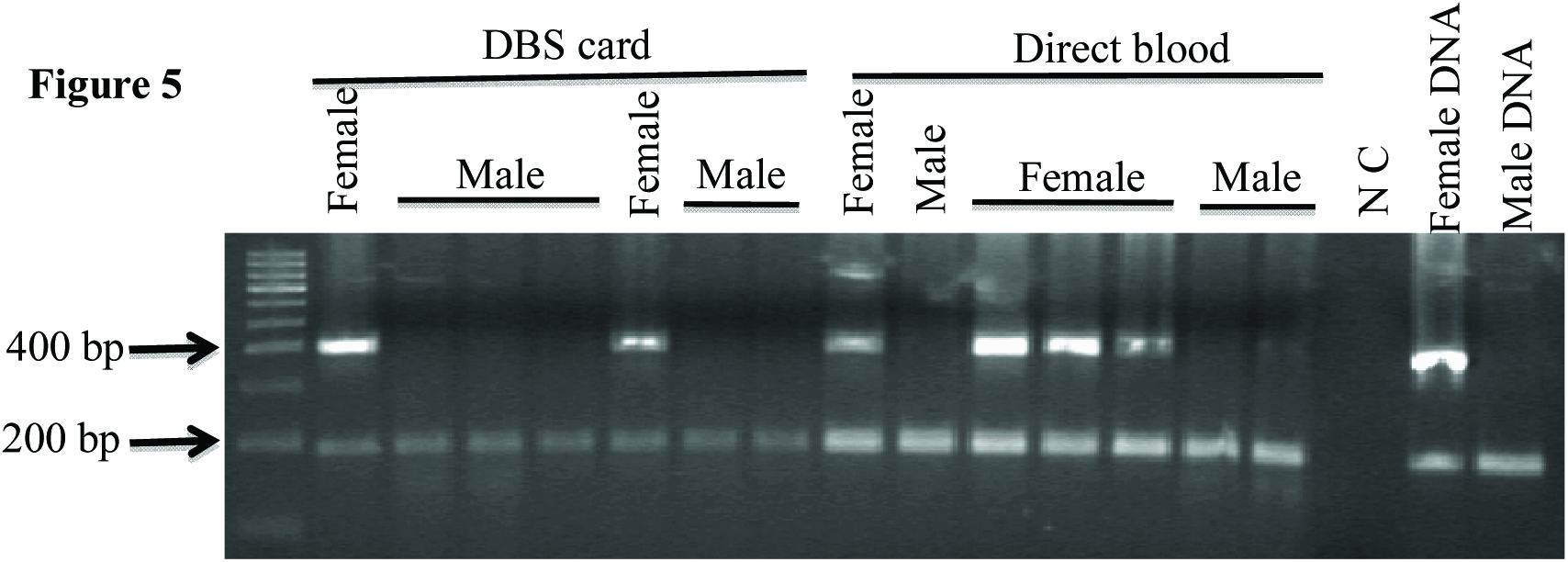
Gender identification of chicken (*Gallus gallus*) by PCR amplification of CHW2F/2R and 18s rRNA gene sequences from whole blood and DBS card. The gene targets was amplified from EDTA treated blood samples obtained from chicken as template. Taq DNA polymerase (5 U) and 1μl of 200 μM each forward and reverse primer were added to a total reaction volume of 50 μl. Genomic DNA isolated from male and female blood samples was used as positive control. Negative control was included in without the genomic DNA.

### Gender identification of chicken using Dried Blood Spot (DBS) cards as a template

In our field level blood collection, the DBS on filter paper was found to have several technical advantages than collection of blood in tubes with anti-coagulant. We optimized the blood collection on filter paper and conventional PCR amplification conditions for the economically important avian species. In our observation with chicken blood, approximately 5-10 μl of blood specimen is highly sufficient to perform nearly 5 PCR analyses. The chicken gender was determined using DBS punches by duplex PCR with CHW1FP/1RP and CLD 18s 1FP/1RP or CHW2FP/2RP and CLD 18s 2FP/2RP primer combinations.

In chicken, the CHW1FP/1RP and CLD 18s 1FP/1RP primer combinations produced only female specific and endogenous control with 3 mM MgCl_2_ and 5 units of Taq enzyme (Fig. 3). While the CHW2FP/2RP and CLD 18s 2FP/2RP primer combinations produced a nonspecific amplification of about 800bp, the amplification intensity of targets comparatively less than direct blood or genomic DNA (Fig. 5) with the same volume of MgCl2 and Taq enzyme. Alterations in MgCl2 and Taq enzyme levels lead to either reduced PCR product or multiple non-specific products.

**Fig. 3.**
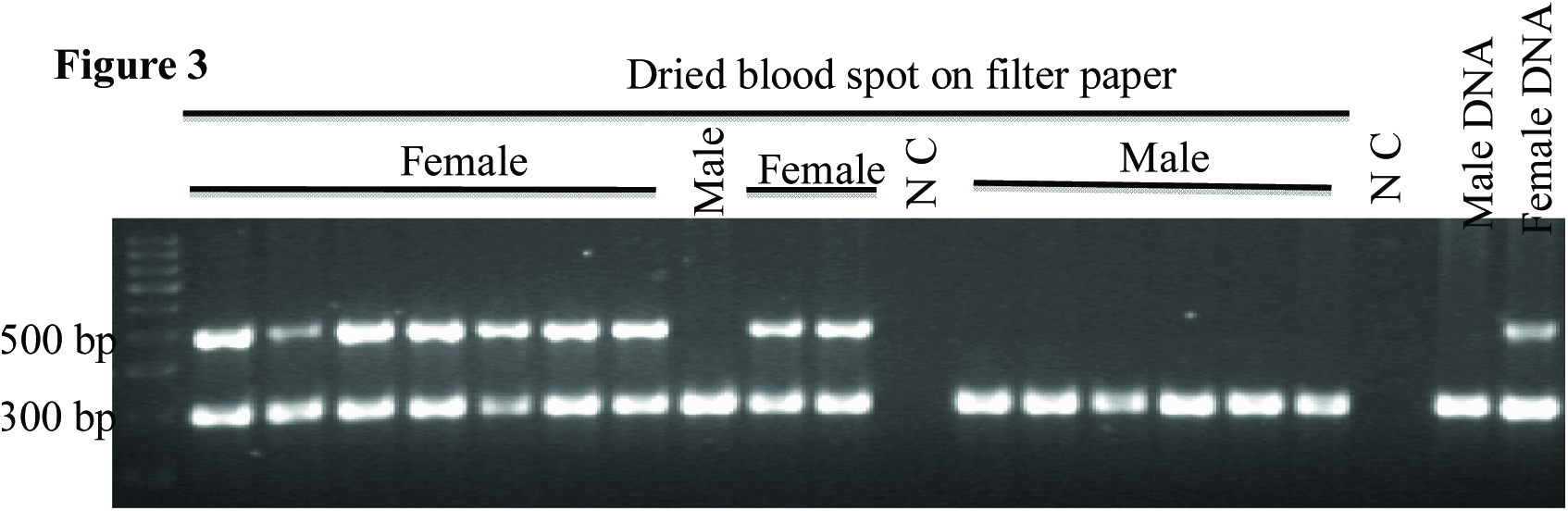
Gender identification of chicken (Gallus gallus) by PCR amplification of CHW1 sequences from dried blood spot (DBS) prepared from male (n=7) and female blood samples (n=9) as the source of template. Taq DNA polymerase (5 U) and 1μl of 150 μM each forward and reverse primer were added to a total reaction volume of 50 μl. Genomic DNA isolated from male and female birds was used as positive control. Negative control was included in each reaction without the genomic DNA.

**Fig. 4.**
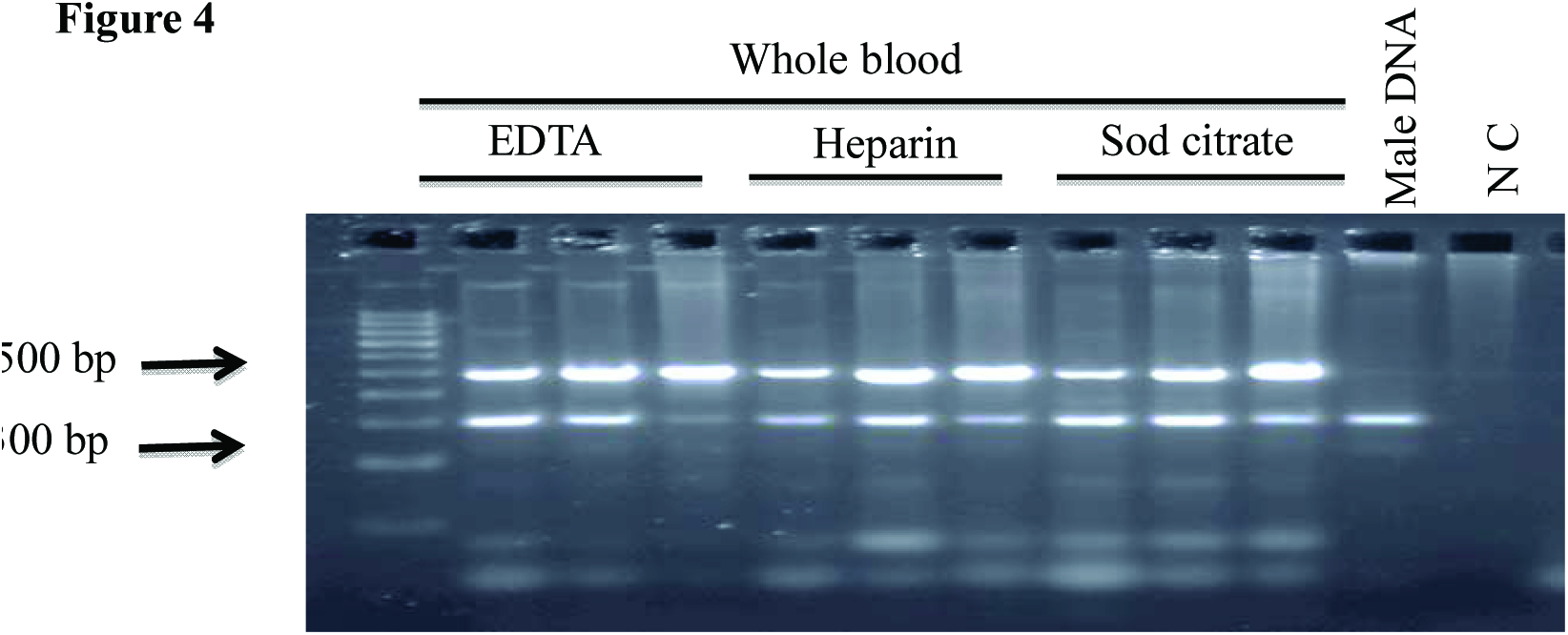
Effect of various anticoagulant treated whole blood as a template in gender identification of chicken (*Gallus gallus*) by PCR amplification of CHW1 and 18s rRNA gene sequences from whole blood. The gene targets was amplified from EDTA, heparin and sodium citrate treated blood samples obtained from chicken as template. Taq DNA polymerase (5 U) and 1μl of 200 μM each forward and reverse primer were added to a total reaction volume of 50 μl. Each well was loaded with 15 μl of PCR products. Genomic DNA isolated from male was used as PCR control. Negative control was included in without the genomic DNA.

**Fig. 6.**
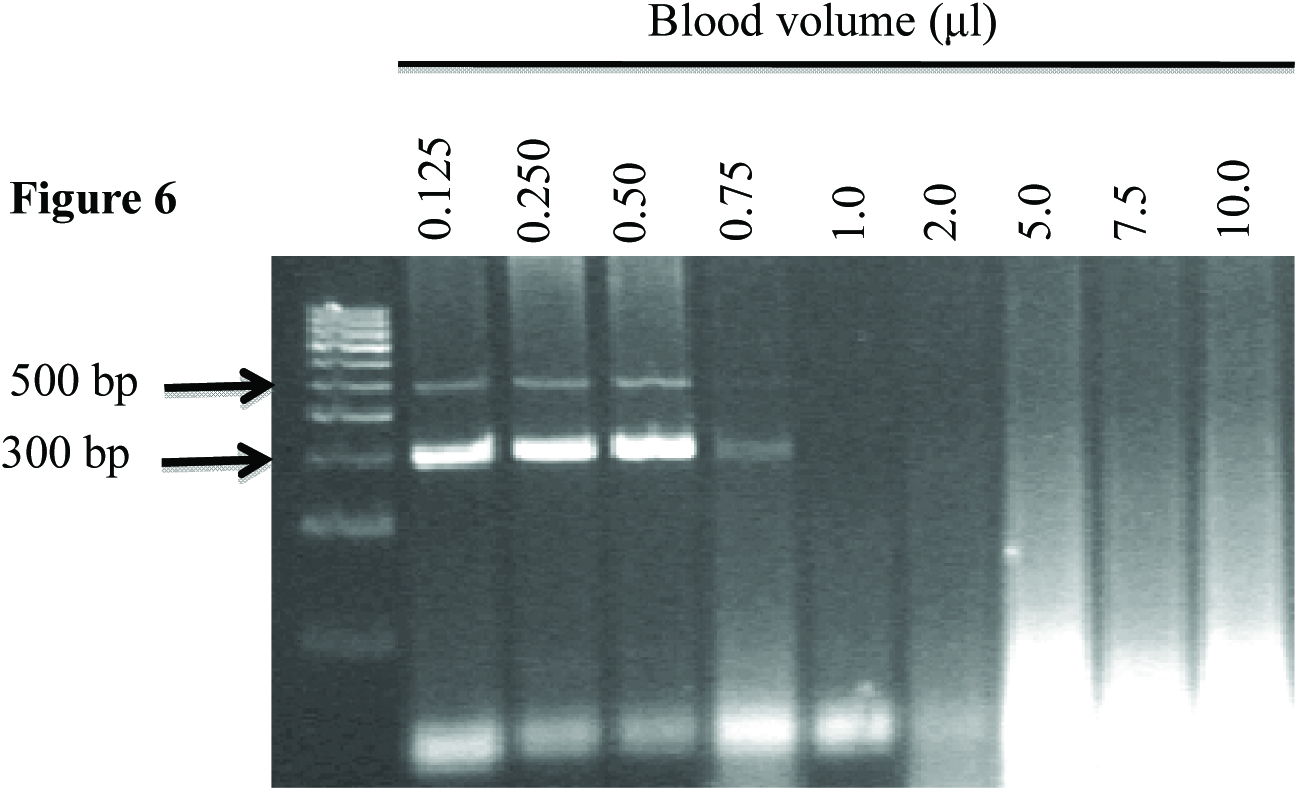
Effect of various volumes of direct blood in PCR amplification of CHW2F/2R and 18s rRNA gene sequences from whole blood and DBS card. The amounts of whole blood used were 0.125, 0.25, 0.5, 0.75, 1, 2, 5, 7.5 and 10 μl. Taq DNA polymerase (5 U) and 1μl of 250 μM each forward and reverse primer were added to a total reaction volume of 50 μl.

## DISCUSSION

PCR based gender determination in avian species is a highly reliable approach in breeding industry, evolutionary and behaviour studies, endangered species conservation and forensic investigations (Freed et al., 2009). PCR based techniques has many advantages for sex identification in sexually monomorphic birds over manual and other methods like ultrasonography, infrared spectroscopic imaging, laparoscopic method, cytogenetic method, hormonal assay (Granevitze et al. 2007, Chen et al. 2012, Lee et al. 2010). Most of the PCR based sex determination methods targets mainly the genes of sex chromosomes. Unlike mammals, the females of avian species are heterogametic, with two different sex chromosomes (ZW), while the males are the homogametic sex (ZZ) (Stevens, L. 1997).

Several studies reported valuable candidate genes for sex determination including CHD1 (chromodomain helicase DNA binding protein 1), HINT (Histidine triad nucleotide binding protein W), ASW (avian sex-specific W-linked gene) and FET1 -(female expressed transcription 1) ATP5A1 (ATP synthase α-subunit), DMRT1 (Double sex and Mab-3-related transcription factor 1), a Z-linked gene supporting the Z-dosage of avian sex determination (Fridolfsson et al. 1998, Stiglec et al. 2007). The feasibility of two chicken W chromosome linked EST genes identified in early female embryos by female-minus-male subtracted cDNA macroarray in sex determination in chicken gained our interest. Surprisingly, the primer pairs designed in this study amplifies only the chicken W-chromosome specific target, which even fail to amplify the female specific targets in evolutionarily related species studied.

Reports on PCR based gender determination approaches and sex-linked markers in avian species are steadily increasing. A continuous effort was made in PCR-based sex determination techniques to increase the accuracy, speed and high-throughput applicability is evident through the previous reports (Morinha et al. 2012). Although PCR based gender determination techniques have several advantages over other techniques, always it demands purified DNA template without any PCR inhibitors for optimal result (Schrader et al. 2012). Addition of whole blood as a template in PCR may influence the amplification by the quantities of heme, lactoferrin and immunoglobin in the reaction mixture. Although many previous reports describes the successful amplification of direct human blood demands pre optimization of PCR buffer, RBC lysis with cell lysis reagents and heat treatment (Nishimura et al. 2000, Yang et al. 2007, Rosenthal et al. 2010). Unlike mammalian blood, nucleated red blood cells and different plasma levels of avian species requires a separate PCR optimisation for the successful amplification of target’s. Different amounts of anticoagulant-treated human blood ranging 0.05−5 μl indicated that 1 ul of any type of anticoagulant-treated blood is enough for a successful 50μl PCR reaction (Bu et al. 2008, Sharma et al. 2012). However the use of 1μl of chicken blood hinders the amplification of target. Our study recommends the use of reduced volume of chicken blood samples ranging from 0.125μl and 0.250μl of in PCR reaction mixture for satisfactory amplification.

Apart from the total volume of blood volume in PCR reaction, the final concentration of MgCl_2_ and Taq polymerase enzyme also plays a significant role in the intensities of target amplification. Although human blood and other biological samples can be directly amplified using 1.5 mM MgCl2 and 1- 2.5 unit of Taq enzyme (Burckhardt 1994, Bu et al. 2008, Li et al. 2011), satisfactory amplification of specific targets using chicken blood samples requires 3 mM MgCl_2_ and 5 units of Taq enzyme. In our observation, 3 mM MgCl_2_ and 5 units of Taq enzyme in a 50μl volume reaction is optimum for the amplification of 3 targets and more than 3 mM MgCl_2_ and 5 units of Taq enzyme affects the intensities of PCR targets. Also we observed that a quick centrifugation after lysis at high temperature results a reasonable PCR yield, this is agreed with the previous report (Bu et al. 2008). Although a previous report describes the amplification of target using whole chicken blood, following pre-treatment of blood samples with NaOH and neutralisation with suitable buffer or pre boiling for the amplification. In our hand, the NaOH treatment hinders with the PCR yield or completely inhibits the PCR reaction in certain samples. Also we observed that the PCR must be carried out with at least 50μl of total reaction volume for successful amplification of chicken blood samples, this contradicts with the previous report which is carried out with 25μl total reaction. (Khatib and Gruenbaum, 1996).

A common problem in collecting blood samples for DNA isolation is preservation of DNA from degradation in the field and during transportation to the laboratory. Although the DNA degradation usually can be prevented through cold chain management, which requires dedicated chilling facilities. The DBS on FTA card is an ideal method when the cold chain management cannot be maintained after sample collection and prior to extraction at field level (McEwen and Reilly 1994, Parker and Cubitt, 1999). However, the use of FTA card and other commercial DNA purification buffers in large scale applications is not feasible due to cost issues (Zhou et al. 2006). Hence we attempted DBS on filter paper for sex determination of avian species, we experienced in this study that this method favours the collection of peripheral blood in micro litres through gentle prick is relatively painless particularly for young birds, easy shipment, minimum involvement of highly skilled personnel, blood spots further represent a low infectious hazard compared to transportation of whole avian blood or biological fluids.

Pre-treatment of FTA card and filter paper containing sheep or human blood samples with NaOH, NaCl and methanol found to be efficient for the amplification of target DNA following DNA extraction (Mc Cabe. 1991, Zhou et al. 2006). However it is necessary to remove any residual levels of NaOH, NaCl and methanol before PCR reaction to achieve optimum amplification. Although many reports suggest the use of commercially available FTA^®^ cards for archiving blood or biological specimens for molecular studies which is not feasible due to the high cost of operation in large scale applications and the chemicals which is used to treat the FTA^®^ card to lyse the cells and to control the pathogens in blood samples and growth of bacteria. The chemicals in the FTA^®^ card require 2-3 hours of purification prior to PCR to reduce the inhibitory effects (Wang et al. 2009). In this study we used filter paper to collect the avian blood samples relatively less volumes (1-5 μl) and ~1 mm of disc was used as template for PCR.

In our observation with direct PCR of chicken blood samples on filter paper, successful amplification of desired target mainly depends on the size of paper punches and volume of blood on filter paper. It is well known that avian blood has nucleated erythrocytes, DBS on filter paper prepared from blood of avian, fish and reptiles results too much DNA for even with small punches to be used directly in PCR reactions (Smith and Burgoyne, 2004). Hence, we applied 1-5 μl either direct or anti-coagulant treated chicken blood on filter paper and reduced the disc size in to approximately 1mm in size. On the other hand, we increased the total PCR volume of 50 μl containing 3 mM MgCl_2_ and 5 units of Taq enzyme to get the optimal amplification.

Over recent years, application of PCR based techniques has great impact in breeding, conservation of endangered species, sex identification of birds and other genetic studies. However, it requires purified DNA from either blood or feather sample whose collection is minimally invasive. Direct PCR from whole blood or dried blood spots will further save time by avoiding DNA isolation. Overall, we consider this method can successfully used for in the gender discrimination and other molecular studies.

## ACKNOWLEDGEMENTS

The author’s greatly acknowledge the Tamil nadu Veterinary and Animal Science University and the Director, centre for animal production for the facilities provided.

